# Mechanistic model of temperature influence on flowering through whole-plant accumulation of *FT*

**DOI:** 10.1101/267104

**Authors:** Hannah A. Kinmonth-Schultz, Melissa J. MacEwen, Daniel D. Seaton, Andrew J. Millar, Takato Imaizumi, Soo-Hyung Kim

## Abstract

We assessed temperature influence on flowering by incorporating temperature-responsive flowering mechanisms across developmental age into an existing model. Temperature influences both the leaf production rate and expression of *FLOWERING LOCUS T* (*FT*), a photoperiodic flowering regulator, in leaves. The *Arabidopsis* Framework Model incorporated temperature influence on leaf growth but ignored the consequences of leaf growth on and direct temperature influence of *FT* expression. We measured *FT* production in differently aged leaves and modified the model, adding the mechanistic temperature influence on *FT* transcription, and linking *FT* to leaf growth. Our simulations suggest that in long days, the developmental timing (leaf number) at which the reproductive transition occurs is influenced by day length and temperature through *FT*, while temperature influences the rate of leaf production and the time (in days) the transition occurs. Further, we demonstrated that *FT* is mainly produced in the first 10 leaves in the Columbia ecotype, and that *FT* accumulation alone cannot explain flowering in conditions in which flowering is delayed. Our simulations supported our hypotheses that: 1) temperature regulation of *FT*, accumulated with leaf growth, is a component of thermal time, and 2) incorporating mechanistic temperature regulation of *FT* can improve model predictions in fluctuating temperatures.

## Introduction

Ambient temperature during the growing season correlates with the timing of plants’ transition from vegetative to reproductive growth. Germination, organ emergence, leaf expansion, photosynthesis, and respiration display similar relationships (Parent *et al.*, 2010). These findings have led to the concept of “thermal time” (Lehenbauer, 1914), a metric that asserts that temperature-driven metabolic rates govern development (Zavalloni et al., 2006), and to models that use the empirical relationship between temperature and development to predict plant response (e.g., Chuine, 2000; Jones *et al.*, 2003).

Thermal time accumulation describes an aggregate of underlying plant responses. Thermal units accumulate more quickly, and reach a predetermined threshold sooner to predict flowering, during warm growing seasons than cool ones. Thermal time implies 1) that all plant physiological rates increase in tandem with temperature increases and 2) that fluctuating and constant temperatures have the same influence on most physiological rates if the mean temperature remains stable. However, processes do not always slow under cool temperatures. The up-regulation of cryo-protective genes (Jaglo-Ottosen *et al.*, 1998) and the circadian clock’s buffering to temperature changes (Rensing & Ruoff, 2002) are just two examples.

Furthermore, predicting plant response to future climates remains imprecise when considering temperature alone or in conjunction with CO_2_ (Asseng *et al.*, 2013; Makowski *et al.*, 2015). The effect of non-stressing temperatures varies among cultivars (Karsai *et al.*, 2013), and plants may respond differently to temperature fluctuations than predicted from constant temperatures (Yin & Kropff, 1996; Kim *et al.*, 2007; Karsai *et al.*, 2008). As most plant models incorporate some variant of thermal time (Ritchie & Otter, 1985; Jamieson *et al.*, 1998a,b; Wilczek *et al.*, 2009; He *et al.*, 2012; Kumudini *et al.*, 2014), they may fail to capture aspects of temperature response. Differing day-length or climate responses may also confound model prediction, with the same cultivar showing different thermal-time requirements, depending on planting date, location, or growth conditions (Piper *et al.*, 1996; Kumudini *et al.*, 2014; Carter *et al.*, 2017). Incorporating the molecular mechanisms of cultivar response in different environments should improve models’ predictive capacity.

A more mechanistic approach would decompose environmental influences into separate model processes (Welch *et al.*, 2003; Kim *et al.*, 2012; Zheng *et al.*, 2013; Brown *et al.*, 2013). One such approach, in wheat, noted that the number of leaves produced before the reproductive transition decreased as the environmental signal’s strength increased (Jamieson *et al.*, 1998b). Prolonged cold, vernalizing temperatures followed by longer days reduced the leaf number at which the transition occurred, while ambient temperature influenced the rate the leaves were produced (Brown *et al.*, 2013). Modeling accumulation of *VRN3*, a key flowering gene, in response to vernalization and day length cues, and as a function of thermal time, accurately predicted final leaf number and timing of flowering (Brown *et al.*, 2013).

*VRN3* is an orthologue of *FLOWERING LOCUS T* (*FT*) in *Arabidopsis thaliana* (Yan *et al.*, 2006), an integrator of environmental cues in the photoperiodic flowering pathway (Song *et al.*, 2015). *FT* levels correlate strongly with the leaf number present when flowering occurs (Krzymuski *et al.*, 2015; Seaton *et al.*, 2015; Kinmonth-Schultz *et al.*, 2016). In turn, day length, vernalization, and ambient temperature changes regulate *FT* expression (Blazquez *et al.*, 2003; Amasino, 2010; Song *et al.*, 2015). *FT* simulated as a function of day length and accumulated as a function of thermal time can accurately predict flowering in some conditions (Chew *et al.*, 2012). Under constant and fluctuating temperature conditions, cool temperatures suppress *FT* through the interaction of SHORT VETIGATIVE PHASE (SVP) and the FLOWERING LOCUS M (FLM)-β splice variant on the *FT* regulatory regions (Blazquez *et al.*, 2003; Posé *et al.*, 2013; Lee *et al.*, 2013; Lutz *et al.*, 2015; Sureshkumar *et al.*, 2016; Kinmonth-Schultz *et al.*, 2016). However, temperature fluctuations from warm to cool induce *FT* through induction of *CONSTANS* (*CO)*, a chief transcriptional activator of *FT* (Schwartz *et al.*, 2009; Kinmonth-Schultz *et al.*, 2016). As there is no simple correlation between temperature decrease and *FT* level reduction, the linear accumulation of flowering gene products with thermal time may not adequately capture the influence of temperature on final leaf number, especially in fluctuating temperatures.

Further, FT protein is expressed in the leaves and moves to the shoot apex where it complexes with FLOWERING LOCUS D (FD) protein to induce the transition from leaf to floral production (Abe *et al.*, 2005; Corbesier *et al.*, 2007). The amount of FT protein perceived at the shoot apex likely depends on the amount of leaf tissue present. Leaf production and growth are strongly temperature dependent (Parent *et al.*, 2010). We proposed that a key mechanism underlying thermal time accumulation could be either the accumulation of gene product (e.g., FT protein) or the increasing capacity for transcript production as a plant grows. In either case, the rate of FT accumulation would be further adjusted by day length and by direct temperature influence on *FT* gene expression.

To predict whole-plant *FT* accumulation we must consider changes in *FT* expression with developmental age. Likely, *FT* expression is not consistent in all leaves or developmental stages. The transcriptional reporter, *pFT:GUS* was expressed in the tips of the two true leaves in six-day-old seedlings, but ranged across the leaf in 12-day-old plants having five to seven true leaves (Takada & Goto, 2003). Further, whole-plant transcript levels increase from age five to 15 days relative to an internal control, indicating changing capacity for *FT* expression with age (Mathieu *et al.*, 2009). *FT* transcript levels have neither been measured in leaves of different ages, nor has this been considered in flowering models, but it could improve our understanding of how day length and temperature impact *FT* to control flowering across developmental age.

In earlier work we found *FT* levels correlate with flowering across a range of temperature conditions ((Kinmonth-Schultz *et al.*, 2016). We also observed that *FT* can be both induced and suppressed by cool temperatures depending on whether constant or fluctuating temperatures are applied (Kinmonth-Schultz *et al.*, 2016). This provided us with an opportunity to determine the relative influences of *FT* transcriptional control versus whole-plant *FT* accumulation via leaf production. One possibility is that despite *FT* induction by a temperature drop, flowering would be delayed because whole-plant leaf production is slowed. Alternatively, *FT* induction could result in flowering times that are earlier than predicted. To address these questions we utilized an existing model (The *Arabidopsis* Framework Model; FM-v1.0) capable of simulating plant growth and flowering times in response to temperature (Chew *et al.*, 2014). We assessed FM-v1.0’s capacity to simulate growth in fluctuating temperature conditions. We then quantified the level of *FT* produced in leaves of different ages and built new models describing the behavior of *FT* across leaves and the influence of fluctuating temperatures on *FT.* We integrated these models into FM-v1.0, linking *FT* accumulation to leaf tissue production. Using this altered model, referred to as FM-v1.5, we explored the sensitivity of *FT* accumulation to both gene expression and leaf growth, and demonstrated how each component may influence flowering times. FM-v1.0 used a more traditional thermal-time approach to determine flowering times, whereas FM-v1.5 uses a more mechanistic approach based on *FT* levels, hence we also compared mechanistic and thermal-time methods of simulating temperature influence on flowering.

## Material and Methods

### Description of Arabidopsis Framework Model and Modifications

The *Arabidopsis* Framework Model (FM-v1.0, Figure S1, Chew *et al.*, 2014) combines plant growth and mechanistic flowering regulation for *Arabidopsis*. FM-v1.0 is run in two phases. In phase one, the timing of flowering is determined by thermal time accumulation (*T(t) − T*_*base*_, calculated hourly) in the Phenology module, with daytime temperature given more weight (Wilczek *et al.*, 2009; Chew *et al.*, 2012). Thermal time is modified by day length, to produce Modified Photothermal Units (MPTUs), through mechanistic circadian- and day-length *FT* transcriptional regulation in the Photoperiodism module (Salazar *et al.*, 2009). The number of days required to reach the MPTU threshold determines the stopping point of vegetative growth and onset of flowering, and is used as an input in phase two. In phase two, the climate parameters affect vegetative growth. Growth is determined by the rate of photosynthesis and carbon partitioning between roots and shoots (Carbon Dynamic module, Rasse & Tocquin, 2006), and includes the rate of organ production as a function of thermal time, including production of individual leaves (Functional Structural Plant module, Christophe *et al.*, 2008). To modify FM-v1.0, we removed the thermal time accumulation used in phase one of FM-v1.0 and instead incorporated mechanistic temperature influence on *FT* into the Photoperiodism module. We maintained thermal time control over leaf tissue production in phase two, but modified the SLA and respiration components to improve the response of leaf growth to fluctuating temperatures. Then, rather than running the model in two phases, we called the Phenology and Photoperiodism modules at each time step, considering their outputs *FT* gene expression per unit of leaf tissue. We used the leaf number, age, and area outputs at each time step to determine the relative *FT* produced by each leaf, and summed the value of *FT* across all leaves to get a whole-plant *FT* value. Our modifications (FM-v1.5, Figure 1) are described in detail below.

**Figure 1.**
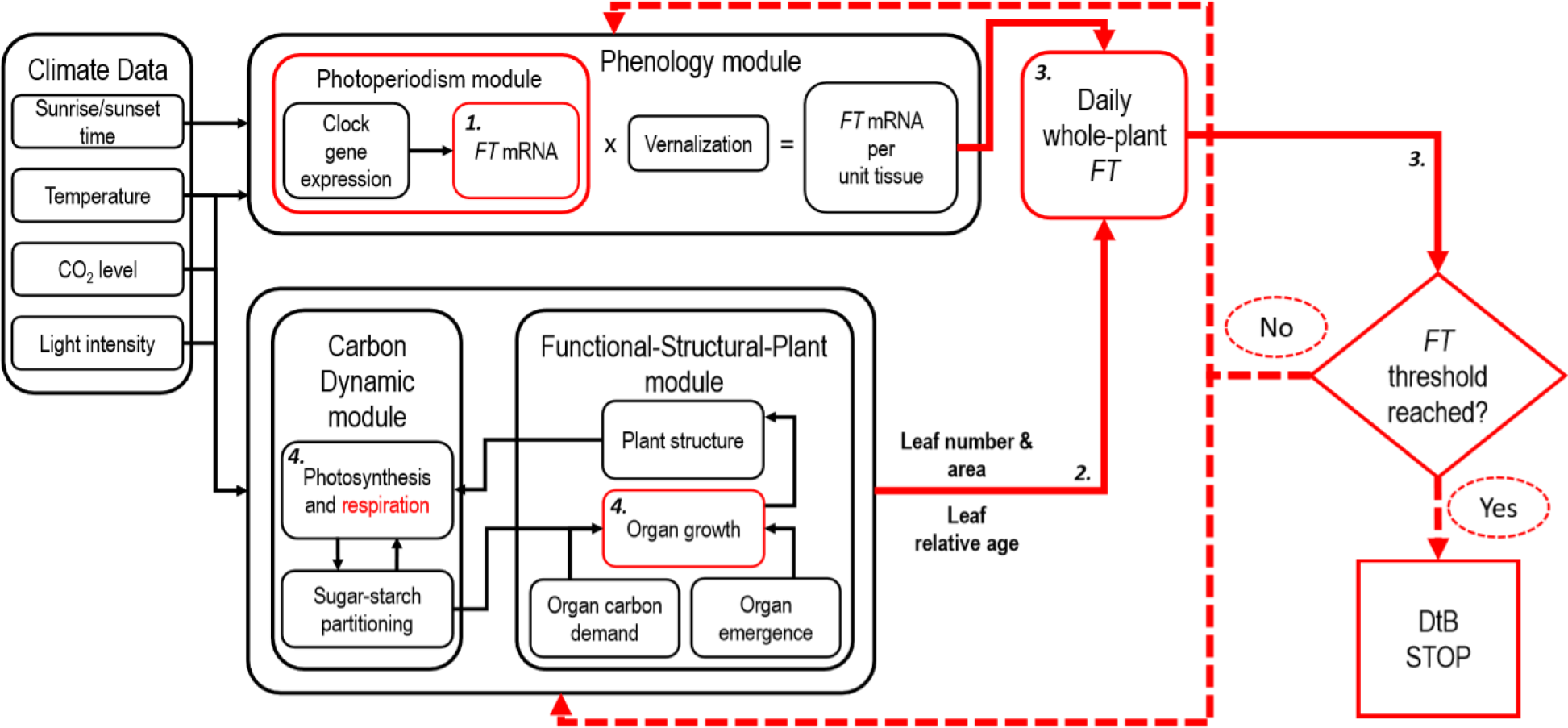
Schematic of Model FM-v1.5. Temperature (through *CONSTANS* and *SHORT VEGETATIVE GROWTH/FLOWERING LOCUSM*), day length, and the circadian clock regulate expression of *FLOWERING LOCUS T* (*FT*) in the Photoperiodism and Phenology modules per unit tissue. The leaf number and relative leaf age, outputs of the Functional Structural Plant module, are used to determine the capacity of each leaf to express *FT*, and leaf area is used to determine the amount of leaf tissue present. *FT* is summed across all leaves in a plant and added to the whole-plant *FT* from the previous time step. The model ceases leaf production and determines the days to bolt (DtB) when *FT* reaches a pre-set threshold set by using the leaf number for plants grown in long days at 22 °C. Red illustrates where adjustments were made to the original model (FM-v1.0). The bold, italic numerals correspond to the numbers in the model description in the main text.

### 1. FT *transcript accumulation in fluctuating temperatures simulated through* SVP *and* CO *influence*

Under long days (LD), in 22 °C day, 12 °C night temperature-cycle conditions (22°C /12 °C-night), *FT* was suppressed at dusk compared to 22 °C constant temperatures (22°C-constant) (Kinmonth-Schultz et al. 2016) likely through the action of SVP and the FLM-β splice variant, consistent with prior observations under constant temperatures (Blazquez *et al.*, 2003; Lee *et al.*, 2007, 2013; Posé *et al.*, 2013). SVP protein levels increased shortly after exposure to cool temperatures (Kinmonth-Schultz et al. 2016), as did the ratio of *FLM-β* to *FLMS-δ* splice variants (Posé *et al.*, 2013). FLM-β facilitates SVP binding, and SVP and FLM-β protein levels increase with decreasing temperatures (Lee *et al.*, 2013). Both SVP and FLM-β are present at 23 °C; a transfer from 23 °C to 27 °C resulted in SVP decay that occurred within 12 h (Lee *et al.*, 2013). We used a single term to simulate the combined SVP and FLM-β behavior termed “SVP activity”. Consistent with the observed behavior of these proteins, we modeled SVP activity to increase in response to a decrease in temperature, as shown below.

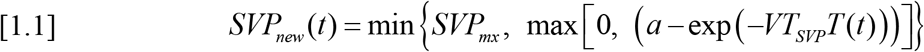

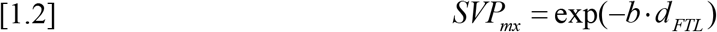

*SVP*_*new*_ is the newly synthesized protein (nmol/h), *VT*_*SVP*_ describes the degree SVP synthesis decreases in response to a temperature increase, the intercept (*a*) is used to adjust the overall amount of SVP synthesized, *T* is temperature (°C), and *t* is time (*t* = 0 at sowing). The influence of SVP may decline over time, as cool-temperature suppression of *FT* disappeared over a two-week period (Figure S2a-b). To simulate this, *SVP*_*mx*_ declines relative to days post emergence of the first true leaves (*d*_*FTL*_, eq. [1.2], Figure S2c). *SVP*_*new*_ is synthesized every hour, and is input into a differential equation calculated continuously [1.3]. Values and units of each coefficient are in Table S1. To capture the suppression of *FT* at dusk, we set the SVP decay rate to be slightly lower than its production. This caused SVP to remain higher at 22 °C after a 12 °C night than in 22°C-constant conditions, even after several hours (Figure S2c). The decay rate (*v*_*SVP*_) is proportional to the present SVP concentration.
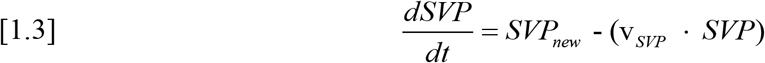

In LD 22°C /12°C-night, *FT* levels are higher at dawn coinciding with higher *CO* mRNA and protein in cool nights (Kinmonth-Schultz *et al.*, 2016). While SVP activity may respond to absolute changes in temperature (Lee *et al.*, 2007, 2013; Posé *et al.*, 2013), *CO* accumulation is induced by rapid changes from warm to cool (Kinmonth-Schultz *et al.*, 2016). The degree of temperature change is likely a factor, as a drop of 10 °C (22°C /12°C-night) yielded more *CO* transcript accumulation than did a drop of 5 °C (22°C/17°C-night) relative to 22 °C constant temperatures (Kinmonth-Schultz *et al.*, 2016). This relationship was linear across the three treatments (Figure S3a). We correlated *CO* mRNA induction (*KT*) linearly with the difference (*dT*) between the maximum and current temperatures (eq. [1.4]). To determine *dT*, the model queries the temperature at each time step, and compares the current temperature against the previous maximum temperature. If higher, the current temperature is set as the new maximum temperature. *dT* may be zero if there has been no decrease in temperature, and *KT* cannot fall below zero.
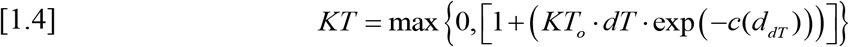

Coefficient *c* describes the rate at which *CO* induction changes with *dT*. The influence of a temperature change fades over several days if the temperature remains cool over that timeframe (Figure S3b). To account for this, *d*_*dT*_ is the time (days) since the change in temperature occurred. *KT* is used to modify the *CO* mRNA (*CO*_*m*_) amount produced (eq. [1.4]), as temperature seems to influence CO through transcription (Kinmonth-Schultz *et al.*, 2016). *CO*_*m*_ is an input for the CO protein (*CO*_*p*_) equation as in Chew *et al.*, 2014, as shown below (eq. [1.5]). Decay occurs only at night (*L*_1_ = Light period).
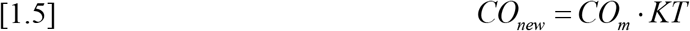

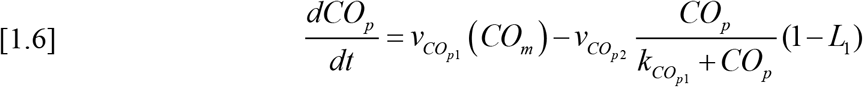

The SVP/FLM-β complex and CO may act competitively at the *FT* promoter (Bratzel & Turck, 2015), with CO overcoming suppression by SVP/FLM-β at night when its levels are high. The Photoperiod module in FM-v1.0 (Chew *et al.*, 2014) describes the relationship between *FT* transcription and CO protein (eq. [S2.1]). We incorporated the interaction between CO and SVP/FLM-β using a modified Michaelis-Menten function for competitive inhibition (Segal, 1976), such that the *k* of *FT* induction by 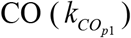 is influenced by SVP activity as below.
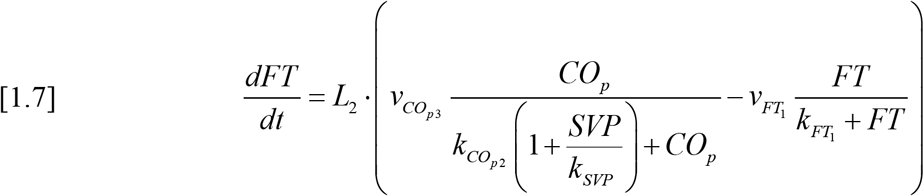

The lower-case *v* and *k* are Michaelis-Menten constants either describing the *FT* synthesis rate as influenced by CO protein (CO_*p*_) or SVP activity, or *FT* degradation. *CO* and *FT* induction were observed when the temperature dropped both at dawn and dusk (Kinmonth-Schultz *et al.*, 2016), like previous observations (Thines *et al.*, 2014). However, daytime *CO* induction was lower than nighttime induction while *FT* induction was higher. The higher daytime CO protein production captured in equation [1.6] was not enough to capture this behavior. While dusk regulation of *FT* is well understood (Song *et al.*, 2015), the mechanisms governing the morning *FT* induction sometimes observed (Corbesier *et al.*, 2007) are not known. To capture the observed behavior, we increased *FT* transcriptional sensitivity in the morning (*L*_*2*_) using a switch function that relied on a model component that peaks around dawn, specifically the circadian clock component, *LHY*, from the Photoperiodism module, (Figure S4). This enabled us to approximate the observed behavior of *FT*.

To entrain the diurnal *FT* and *CO* patterns, we incorporated data from three different treatment types all in 16-h photoperiods grown at ~60 umol m^2^ s^−1^ photon flux density: warm-day (22 °C), cool-night (12 or 17 °C) temperature cycles, in which the temperature change occurred at dusk (24 wild-type replicates, six including 17 °C, and five including the *svp* mutant line); constant warm (22 °C) temperatures shifting to constant cool (12 or 17 °C) temperatures at dawn (eight and three replicates respectively); and growth at 12 and 17 °C from seed (three replicates each) (Kinmonth-Schultz *et al.*, 2016). In all instances, growth from seed at 22 °C was used as the control. The temperature-cycle harvests including 17 °C spanned two days. An ANOVA comparison of models including and excluding *day* as a factor, showed no difference. The days were counted as separate replicates for model training. *FT* and *CO* gene expression were pooled across all replicates within a treatment and normalized across treatments to the mean peak *FT* expression (ZT 16) and mean peak *CO* expression (ZT 16 and 20 mean) in the 22 °C control. Parameter values for change in *CO* induction and *SVP* activity over a period of days were determined using experiments with four replicates each (Kinmonth-Schultz *et al.*, 2016). As we were interested in the cumulative influence of *FT*, we assessed model fit and performance in three ways: (1) minimizing RMSE between observed and predicted gene expression profiles over the 24-h harvest period (14 d after sowing), (2) comparing observed and predicted amounts of *CO* and *FT* as calculated as the area under the curve (AUC) 14 d after sowing, and (3) maintenance of gene expression patterns through time.

### 2. *Incorporating* FT *as a function of leaf and plant age*

We found that *FT* expression declined as leaves aged. Newer leaves in older plants seemed to lose capacity to express *FT* (Figure 2, S5). To simulate the proportion of *FT* per unit tissue (FT, nmol cm^−2^) of each leaf, we used a beta function (eq. [3.1], Yin *et al.*, 1995) based on relative leaf age (*r*), beginning with the youngest emerged leaf as leaf one.
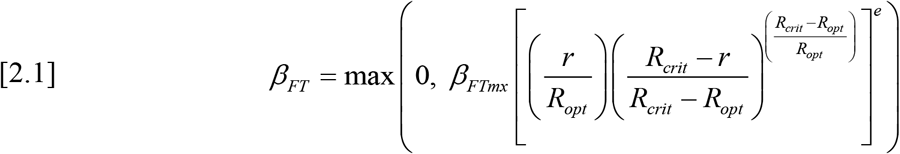

*β*_*FT*_ yields a value between zero and one. *β*_*FTmx*_ describes the maximum value that can be attained by a leaf of a single plant, *R*_*opt*_ is the relative age at which that maximum value is attained, *R*_*crit*_ is the oldest leaf that can express *FT*, and *e* describes the steepness of the curvature. This function causes the dependent variable to oscillate if the independent variable spans a broad range. To avoid this behavior, we set *β*_*FT*_ to be zero below and above the relative ages where *β*_*FT*_ first attains a minimum. *β*_*FTmx*_ and *R*_*opt*_ are dependent on the total number of leaves on a plant (*l*), as described below, avoiding the need to reparameterize for plants of different ages. *f* and *g* are coefficients.
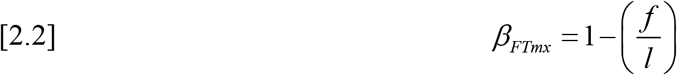

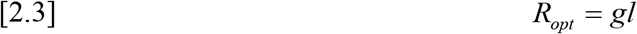

**Figure 2.**
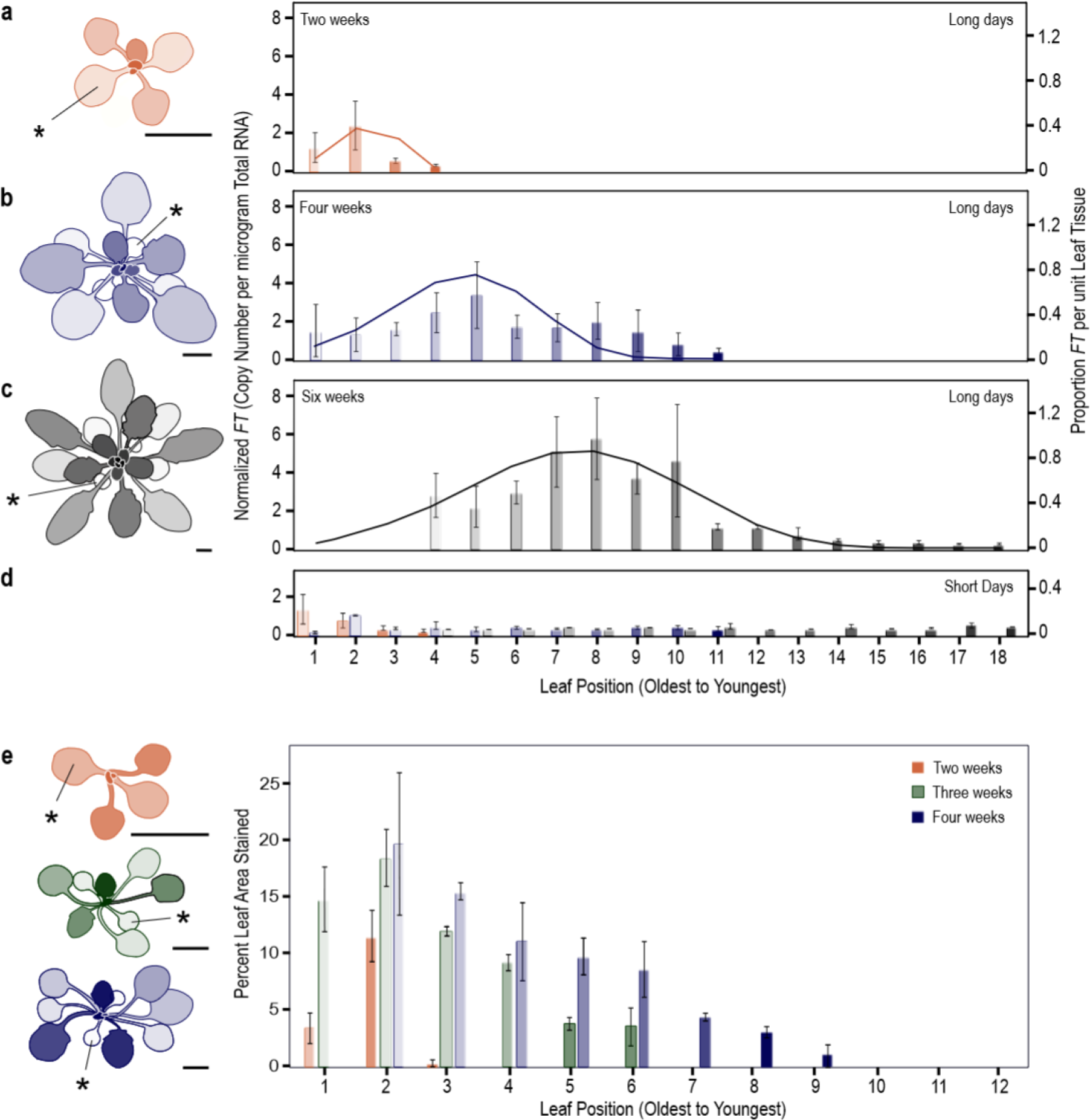
*FT* expression declines in later produced leaves. Leaves of plants aged two **(a)**, four **(b)**, and six **(c)** weeks old and grown in short days were exposed to long days or short days **(d)** for three days, then harvested at 16 hours after dawn on the third day to determine *FT* amount per leaf. The colors in **(d)** correspond to the colors and ages in panels **(a-c)**. *FT* levels were determined by absolute copy number and normalized within a replicate. The simulated proportion of *FT* per unit leaf tissue (cm^−2^, solid lines) for each plant age is shown. This value was used in FM-v1.5 as a modifier to adjust the amount of *FT* produced by each leaf. Percent of the leaf area showing staining in *pFT:GUS* plants **(e)**. For all, the two cotyledons and first two true leaves were pooled for each sample as they emerge in pairs. Older leaves in the six-week old plants failed to yield 2μg total RNA and were excluded. For each plant inset, asterisk indicates one of each cotyledon pair. The shading of the bar graphs (light to dark) indicates leaf age (oldest, first to emerge, to youngest) and corresponds to the shading in the plant insets. Scale bars = 0.5 cm.

### 3. *Determining whole-plant* FT *levels and accumulating* FT *to a threshold*

To link *FT* transcript accumulation to leaf tissue production, the Phenology module is called at each time step. We consider the output of the Phenology module to be the amount of *FT* produced per unit leaf area (*FT*, nmol cm^−2^). This value is adjusted by leaf area (*LA*, cm^2^) and capacity of each leaf to express *FT* (*β*_*FT*_, unitless modifier), as *FT* induction is dependent on light intercepted by the leaf.
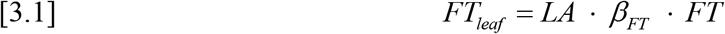

At each time step, *FT*_*leaf*_ (nmol leaf^−1^) is determined for each leaf, summed across all leaves, and added to the value from the previous time step to determine whole-plant *FT* levels. Such *FT* accumulation is consistent with the observation that several days of *FT* induction are needed to induce flowering (Corbesier *et al.*, 2007; Krzymuski *et al.*, 2015; Kinmonth-Schultz *et al.*, 2016). To predict flowering, the model runs until a threshold level of *FT* is reached. This threshold is determined by simulating whole-plant *FT*, at constant 22 °C in LDs, accumulated until a target leaf number is reached. All other treatments are run to this threshold under the assumption that it remains conserved under different growing temperatures.

In FM-v1.0, the development rate towards flowering, as influenced by *FT* amount and photoperiod, is limited below and above two critical daylengths (10 and 14 h) using a different parameter set for each photoperiod (Chew *et al.*, 2014).
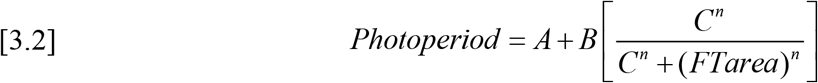

Here, we removed this function and considered direct *FT* accumulation. Determining the absolute amount of *FT* required to induce flowering and whether there are threshold levels of transcription, below and above which flowering time is unaffected, will be a useful future study. We maintained the vernalization component from FM-v1.0 to maintain model flexibility, as vernalization should modify overall levels of *FT* (Helliwell *et al.*, 2006; Searle *et al.*, 2006). This value falls between zero and one and now modifies the levels of *FT* produced within the Phenology model rather than modifying the thermal unit accumulation rate.

### 4. Adjusting FM-v1.0 leaf-area response to fluctuating temperature

FM-v1.0 was parameterized for constant temperatures. It captured the leaf areas of plants exposed to different constant temperatures, but simulated larger areas for plants grown in fluctuating temperatures than the constant-temperature control (Figure S6a). Observed plants accumulated similar biomass, but a lower Specific Leaf Area (SLA, m^2^ g^−1^) under fluctuating temperatures relative to a constant-temperature control (Pyl *et al.*, 2012). In FM-v1.5, we adjusted the SLA and respiration components to improve the relationship among leaf areas across fluctuating temperature conditions (described below).

The larger leaf area under fluctuating temperatures in FM-v1.0 occurred for two reasons. First, SLA decreases with increasing thermal time (i.e. developmental time, Christophe et al. 2008). In FM-v1.0, this causes simulated SLA to be lower in warmer conditions because development is faster (Figure S6b-c), while all treatments begin at a similar biomass. Second, FM-v1.0 relates maintenance respiration to temperature using the Arrhenius function, causing respiration to be lower under cooler temperatures. Under warm daytime temperatures, plants simulated in fluctuating temperatures accumulate the same amount of stored carbon as the control (Figure S6d). Once shifted to cooler temperatures, the lower maintenance respiration rate (Figure S6e) leaves a larger stored carbon pool that can be used for growth, causing larger leaves.

Respiration, carbon storage, or growth may be altered by temperature in ways not captured in the model. In cold-tolerant woody species, respiration of stem cuttings increased near freezing, rather than following the trend predicted by the Arrhenius function, as did the pool of non-structural carbohydrates (NSC) (Sperling *et al.*, 2015). Respiration may also increase at more moderate temperatures in cases where freezing tolerance is induced, as in *Arabidopsis* at 16 °C in light with a low red/far-red ratio (Franklin & Whitelam, 2007). In chrysanthemum, cool nighttime temperatures decreased leaf area while increasing dry weight, by increasing stored starch (Heinsvig Kjær *et al.*, 2007). FM-v1.0 does not incorporate these complexities nor consider sinks for carbon other than growth, such as NSCs. Therefore, to simulate the relative relationships in leaf area across temperature conditions needed for our study (Figure S6f), we removed the temperature sensitivity of maintenance respiration and adjusted the Specific Leaf Area (SLA, m^2^ g^−1^) to decline with decreasing temperature using observations from Pyl et al. 2012 (Figure S7). A more accurate representation of respiration and carbon pools should be incorporated into future models to improve plant growth predictions in a range of temperature conditions.

*Plant growth conditions, RNA expression, GUS tissue analysis, and statistical analysis and experimental controls can be found in the supplemental material*.

## Results

### *Behavior of* CO *and* FT *transcript accumulation in fluctuating temperatures in FM-v1.5*

The *FT* induction by fluctuating temperatures was incorporated through *CO* transcript, which was induced in response to a change to cool temperatures like that observed (Figure 3a-b). There was a strong relationship between the amount of simulated and observed *CO* transcript across treatments, as calculated as the area under the curve (AUC, Figure 3c); although, FM-v1.5 does not incorporate the *CO* suppression observed when plants are grown at constant 12 °C from seed (12°C-constant) (Kinmonth-Schultz *et al.*, 2016). These model modifications, coupled with increased *FT* transcriptional sensitivity near dawn, resulted in induction of *FT* after a temperature drop at dawn or dusk like that observed (Figure 3d-e). Suppression of *FT* through SVP activity, mimicked the observed *FT* suppression at dusk. When the SVP influence is removed in FM-v1.5 to mimic an *svp* mutant, dusk *FT* suppression in warm-day, cool-night conditions (22°C /12°C-night) disappears as is observed; however, simulated morning induction of *FT* is higher, perhaps because SVP activity accounts for both SVP and FLM-β (Figure S8). This strong induction through *CO* was necessary in FM-v1.5 to simulate *FT* induction by cool temperatures in wildtype.

**Figure 3.**
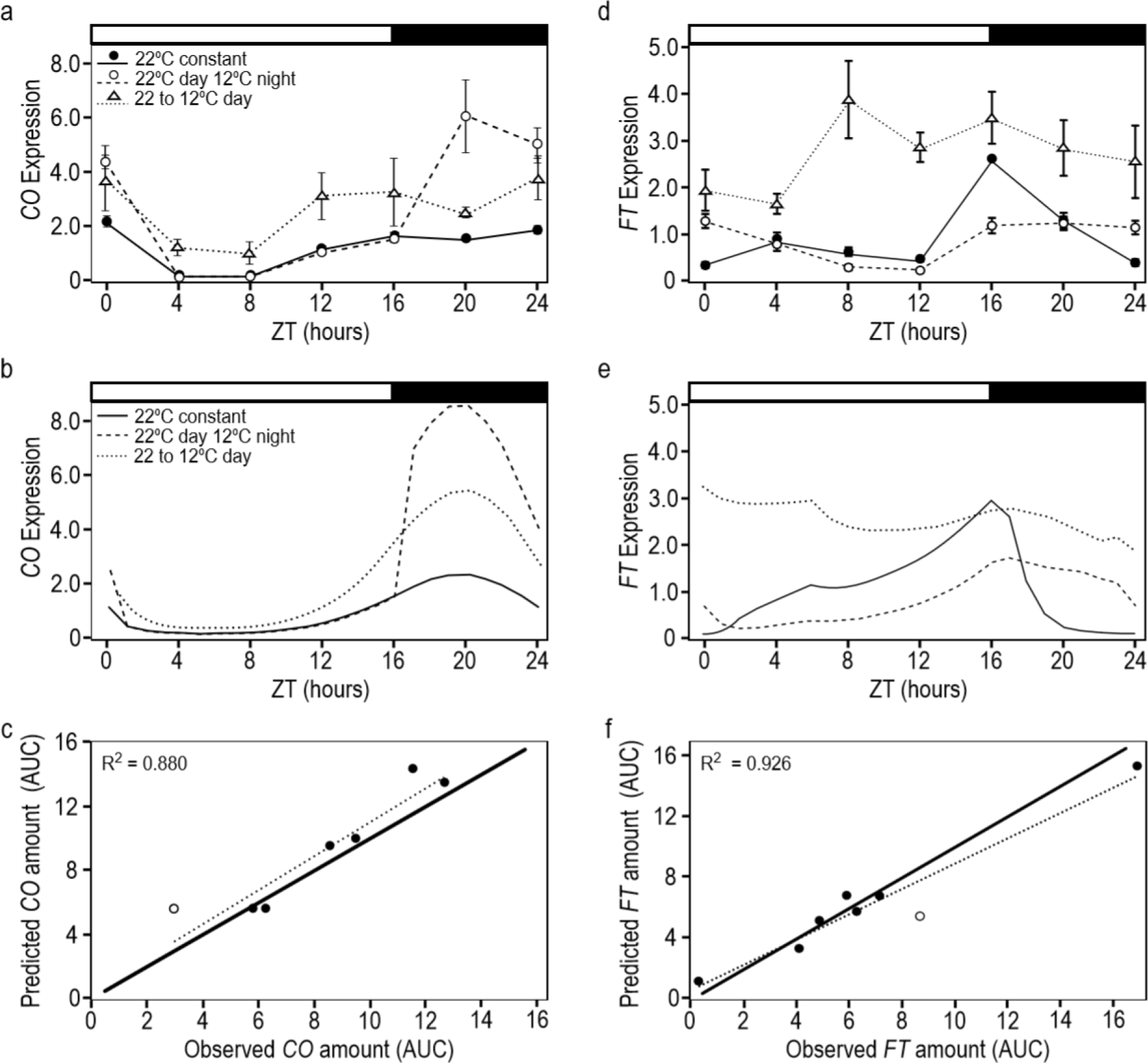
FM-v1.5 mimics general behaviors of *CO* and *FT* in response to temperature, and can accommodate the overall change in amount across treatments. Observed **(a, d)** and predicted **(b, e)** diurnal patterns of *CO* **(a, b)** and *FT* **(d, e)** gene expression in warm (22 °C)-day, cool (12 °C)-night temperature-cycle treatments and in conditions in which the temperature dropped from 22 °C to 12 °C at dawn, then remained at the cooler temperature (22 to 12 °C day) relative to the 22 °C-constant temperature control. The y-axis **(a, b, d, e)** is in zeitgeber time (ZT), and represents hours after dawn. The white and black bars represent light and dark periods respectively. Error bars = 1 S. E. If error bars are not visible, the S. E. is smaller than the height of the symbol. Correlation between predicted and observed results for *CO* **(c)** and *FT* **(f)**, as calculated as the area under the curve (AUC) four days after temperature treatments are imposed. Treatments include warm-day, cool-night cycles, drops to cooler temperatures at dawn, and growth from seed at constant temperatures. All treatment groups include 12, 17 and 22 °C. Dotted lines = correlation, solid lines = one-to-one line. Open circles are growth from seed at 12 °C **(c)** and drop from 22 °C to 17 °C at dawn **(f)**.

For flowering to occur, favorable conditions must occur over several days (Kinmonth-Schultz *et al.*, 2016; Krzymuski *et al.*, 2015; Corbesier *et al.*, 2007). Our aim was to approximate *FT* behavior within a day and through time. Observed *FT* suppression at dusk in 22/12°C-night conditions occurs by day two of the temperature-cycle treatment (Figure S2a). This is true with FM-v1.5 as well, although *FT* levels continue to decline until day four relative to the constant-temperature control (Figure S2b). Over two weeks, the increase in dusk *FT* levels in 22/12°C-night conditions relative to the 22°C-constant control is similar between observed and simulated data (Figure S2a-b). Together, FM-v1.5 can accommodate the wide range in *FT* transcribed across treatments (Figure 2f), and *FT* behaves similarly over time to that observed, allowing us to explore the temperature influence on *FT* expression and flowering in LDs.

### *Assessment of* FT *accumulation in FM-v1.5 across temperatures*

FM-v1.5 allows us to assess the relative temperature influence on *FT* accumulation through both gene expression and leaf development. We compared the total *FT* accumulated 9 days post emergence, equivalent to 1 week in fluctuating temperature treatments, considering 1) influence of temperature on gene expression only (GE), 2) *FT* accumulated with leaf tissue production as influenced by thermal time, temperature influence on gene expression excluded (LTP), and 3) gene expression changes incorporated with leaf tissue production (LTP+GE, full FM-v1.5 model). The influence of age on a leaf’s capacity to express *FT* is incorporated into both the LTP and LTP+GE model variants.

When considering LTP+GE, total *FT* declined, relative to the 22°C-constant control, with increasing exposure times to cool temperature as would be expected from leaf area changes (Figure 4a). When only transcriptional changes were considered (GE), *FT* accumulated at a faster rate than the control for some treatments (i.e. a drop in daytime temperature, Figure 4b). For treatments in which *FT* accumulated more slowly than the control, as in 12°C-constant, the relative difference from the control was less extreme than in LTP+GE. For comparison, we explored the relative difference in accumulated MPTUs, which control flowering time in FM-v1.0, over this timeframe. MPTUs across treatments differed to a lesser degree than accumulated *FT* transcript in LTP+GE, even when nighttime temperatures carried the same weight as daytime temperatures (Figure 4a).

**Figure 4.**
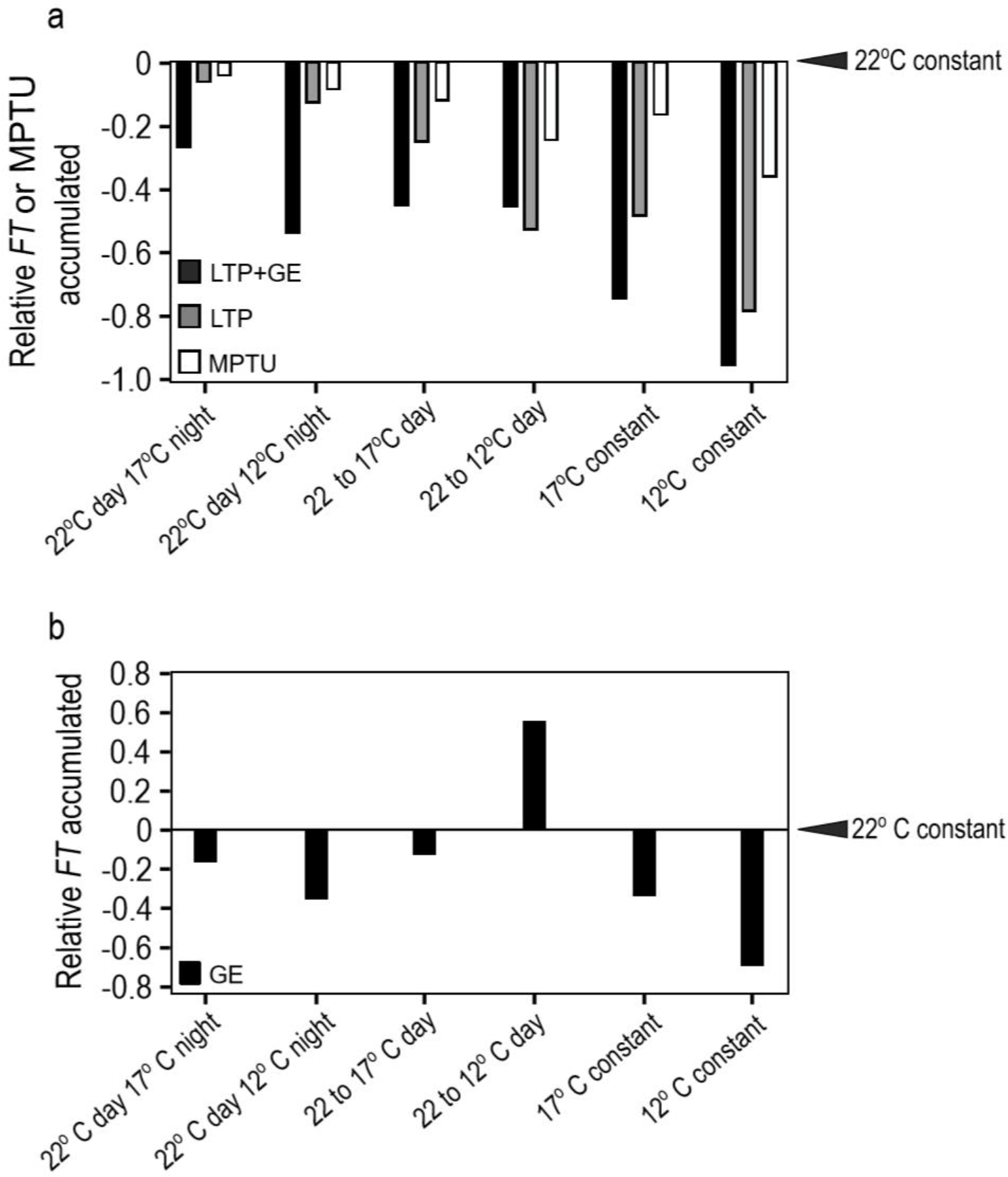
**(a, b)** Whole-plant *FT* accumulation influenced by temperature in fluctuating and constant-cool temperature conditions, differs more strongly from the 22 °C control than does accumulated Modified Photothermal Units (MPTUs). Total *FT* accumulated in constant and fluctuating temperature conditions relative to 22 °C constant temperatures (indicated by arrowheads) 9 ds post emergence, equivalent to 1 wk in fluctuating temperature treatments. **(a) LTP+GE:** *FT* accumulation in full FM-v1.5 model, i.e. temperature affects *FT* gene expression though *CO* and SVP/FLM as well as through leaf tissue production; **LTP:** *FT* accumulation only with leaf tissue production as influenced by thermal time, temperature influence on *FT* gene expression excluded; MPTU: Accumulated Modified Photothermal Units from FM-v1.0. Here, daytime and nighttime temperatures are given equal weight. **(b) GE:** *FT* accumulation considering only influence of temperature on *FT* gene expression, decoupled from leaf production. *22 °C day 12 or 17 °C night* indicates warm-day, cool, night cycles, *22 to 12 or 17 °C day* indicates treatments in which the temperature drop occurred at dawn, then remained cool for the duration of the experiment, *constant* indicates temperatures remained constant from seed.

To assess the influence *FT* transcriptional changes due to temperature have on whole-plant *FT* levels, we used the LTP model variant, meaning that temperature influenced *FT* only through leaf production modulated by thermal time. LTP did differ in whole-plant accumulation. Total *FT* accumulation in the warm-day, cool-night temperature cycle treatments moved closer to that of the control compared to LTP+GE (Figure 4a). When the daytime temperature dropped from 22 °C to 12 °C (22/12°C-day) *FT* accumulated more quickly in LTP+GE than in LTP.

### Assessing capacity of FM-v1.5 to predict flowering

How well can *FT* accumulation predict flowering? What impacts do transcriptional changes have compared to that of whole-pant *FT* accumulation? To assess this, we simulated experiments for plants grown in warm-day, cool-night temperature cycles (Kinmonth-Schultz *et al.*, 2016) as plants often experience such temperature fluctuations in nature. We first assumed that *FT* accumulates to a threshold in a manner like thermal time accumulation. This assumption is consistent with observations that *FT* induction must occur over a period of days before flowering is induced (Corbesier *et al.*, 2007; Krzymuski *et al.*, 2015; Kinmonth-Schultz *et al.*, 2016). We set the threshold as the value of *FT* accumulated when plants reached 15 and 8 leaves, which was the nearest whole number to the average leaf number at bolt for Columbia-0 (Col-0) and Landsburg *erecta* (L*er*), respectively, grown in LD 22°C-constant conditions (Kinmonth-Schultz *et al.*, 2016). We maintained the strain-specific parameters for rate of emergence and leaf initiation from FM-v1.0, as they were comparable to our results (Figure 5a), but added a 7-d delay after initiation of the final leaf to improve the fit across strains at 22 °C. This was to account for the time between initiation of the leaf primordia as modeled (Christophe *et al.*, 2008) and growth of a visible bolt, counted when the stem below the bolt head was 2 mm in length (Kinmonth-Schultz *et al.*, 2016).

**Figure 5:**
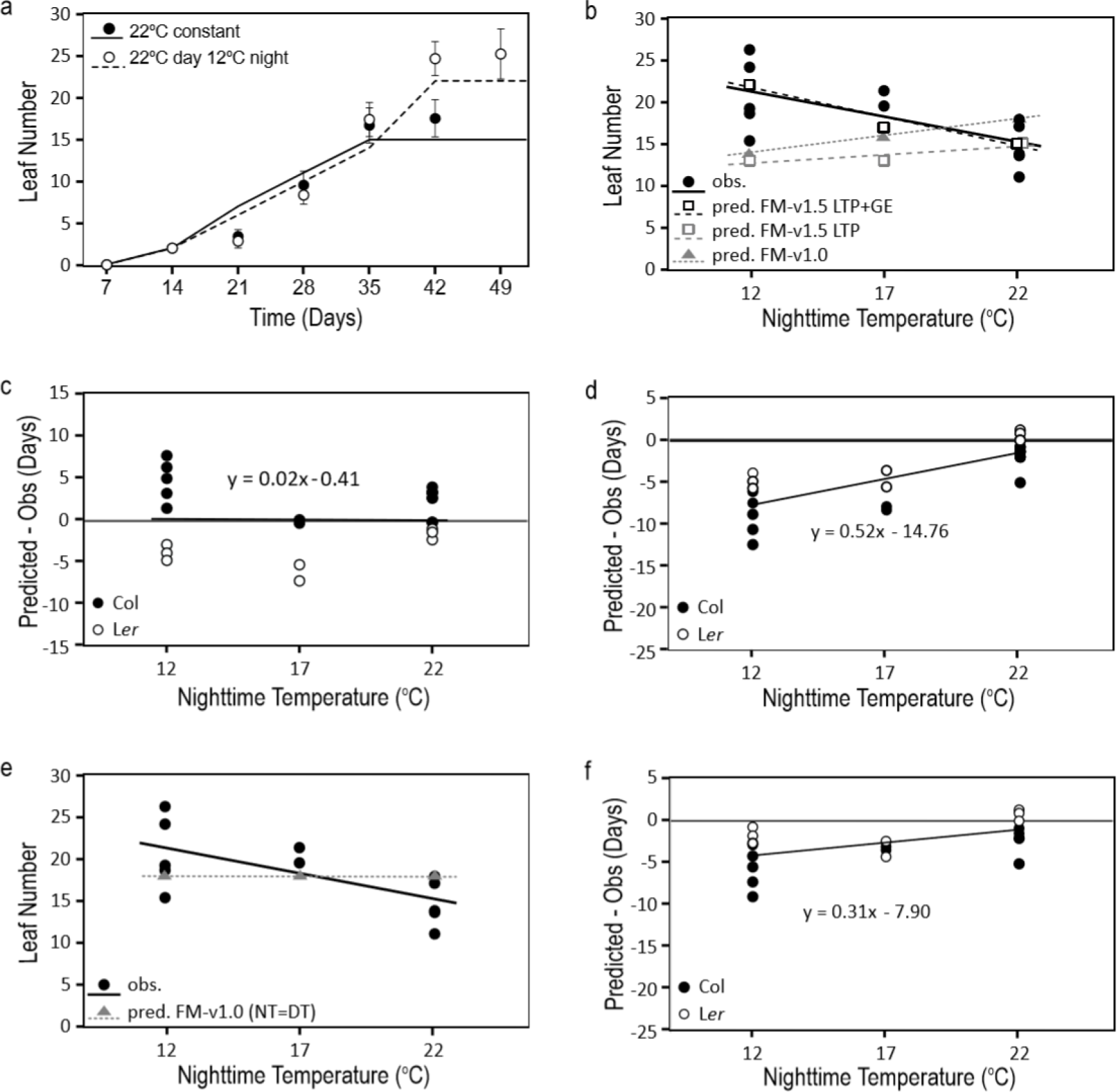
*FT* accumulation as influenced through *CO* and SVP/FLM and leaf tissue production can improve model predictions in fluctuating temperature conditions compared to Modified Photothermal Units (MPTUs). **(a)** Comparison of simulated (lines. FM-v1.5 LTP+GE) and observed (symbols) leaf number by week in Col in constant 22 °C conditions and in 22 °C-day, 12°C-night temperature cycles. **(b)** Final leaf number of Columbia-0 (Col) at bolt as observed (obs.) and predicted (pred.) by incorporating temperature influence on *FT* though leaf tissue production (LTP) and *FT* gene expression (GE) (FM-v1.5 LTP+GE), leaf tissue production only (FM-v1.5 LTP), and through traditional Modified Photothermal Units (MPTU) in FM-v1.0. **(c, d)** The difference between predicted and observed days to bolt in Columbia-0 (Col) and Landsberg *erecta* (L*er*) using FM-v1.5 LTP+GE **(c)** and MPTUs in FM-v1.0 **(d)**. **(e)** Observed and predicted final leaf number and **(f)** the difference between predicted and observed results using MPTUs in FM-v1.0, adjusted so that daytime and nighttime temperatures are given equal weight. **(b-f)** Plotted over three nighttime temperatures. Daytime temperature was 22 °C. **(c, d, f)** Horizontal line at zero is the position in which there is no difference between predicted and observed results. Error bars = 1 S. D. If error bars are not visible, the S. D. is smaller than the height of the symbol.

We then compared the predicted final leaf number and days to bolt for warm-day, cool-night temperature-cycle treatments in the LTP and LTP+GE model variants in FM-v1.5. Cool temperatures delay bolting and increase leaf number (Blazquez *et al.*, 2003, Kinmonth-Schultz *et al.*, 2016). In LTP, we expected that cool-nighttime temperatures would cause flowering to occur at a similar leaf number to the 22°C-constant control because temperature was not influencing gene expression; however, plants would still flower later due to slower whole-plant *FT* accumulation through slower leaf growth. LTP predicted a trend opposite that observed, with a lower leaf number for both 22/17°C-night and 22/12°C-night treatments (Table 1, Figure 5b), because leaves that are present continue to produce *FT* such that it accumulates over time as well as with leaf growth. This caused *FT* to reach the threshold at a lower leaf number. As expected, both 22/17°C-night and 22/12°C-night treatments bolted later than the 22°C-constant control (Table 1). The full LTP+GE variant followed a trend close to that observed, increasing the final leaf number for both cool-night temperature treatments and causing a stronger delay in days to bolt than LTP (Table 1, Figure 5a-c).

**Table 1:** Observed and simulated days to bolt and leaf number in Columbia-0 (Col-0) and Landsberg *erecta* (L*er*) plants exposed to short-term drops in temperature.

| Strain |  | Treatment | Obs. data | FM-v1.0 | FM-v1.5<br>LTP+GE | FM-v1.5<br>LTP |
| --- | --- | --- | --- | --- | --- | --- |
| Col-0 | Days to bolt | 22 °C day 22 °C night | 32.27 | 30.33 | 35.00 | 35.00 |
|  |  | 22 °C day 17 °C night | 38.60 | 30.63 | 38.50 | 35.96 |
|  |  | 22 °C day 12 °C night | 40.08 | 31.25 | 44.88 | 37.50 |
|  | Leaf number | 22 °C day 22 °C night | 14.77 | 18.00 | 15.00 | 15.00 |
|  |  | 22 °C day 17 °C night | 20.50 | 16.00 | 17.00 | 13.00 |
|  |  | 22 °C day 12 °C night | 20.79 | 14.00 | 22.00 | 13.00 |
| Ler | Days to bolt | 22 °C day 22 °C night | 27.13 | 27.75 | 25.42 | 25.42 |
|  |  | 22 °C day 17 °C night | 32.75 | 28.29 | 26.50 | 25.67 |
|  |  | 22 °C day 12 °C night | 33.23 | 28.63 | 29.33 | 25.79 |
|  | Leaf number | 22 °C day 22 °C night | 7.40 | 14.00 | 8.00 | 8.00 |
|  |  | 22 °C day 17 °C night | 8.17 | 13.00 | 8.00 | 7.00 |
|  |  | 22 °C day 12 °C night | 9.98 | 12.00 | 9.00 | 7.00 |

We compared this behavior to flowering predicted using MPTU accumulation by FM-v1.0, adjusting the MPTU threshold to our LD 22°C-constant conditions, as recommended (Chew *et al.*, 2014). If FM-v1.0 adequately captured temperature influence, then the MPTU threshold should be similar across treatments, with negligible differences between predicted and observed results for all three temperature regimes. FM-v1.0 predicted fewer leaves in both 22/12°C-night and 22/17°C-night conditions than in the 22°C-constant control, because it reached the MPTU target before reaching the observed final leaf number (Table 1, Figure 5b). FM-v1.0 accurately captured days to bolt for Col-0 and L*er* grown in 22°C-constant conditions, and showed an expected delay in days to bolt for both 22/12°C-night and 22/17°C-night. However, the days to bolt were lower than observed (Table 1, Figure 5d). Recalibrating to equalize the influence of nighttime and daytime temperatures (daytime temperatures are given more weight in FM-v1.0 (Chew *et al.*, 2012)) reduced but did not eliminate these trends (Table 1-2, Figure 5e-f). Therefore, incorporating mechanistic *FT* accumulation can improve model predictions in fluctuating ambient temperature conditions (Table 2).

**Table 2:**
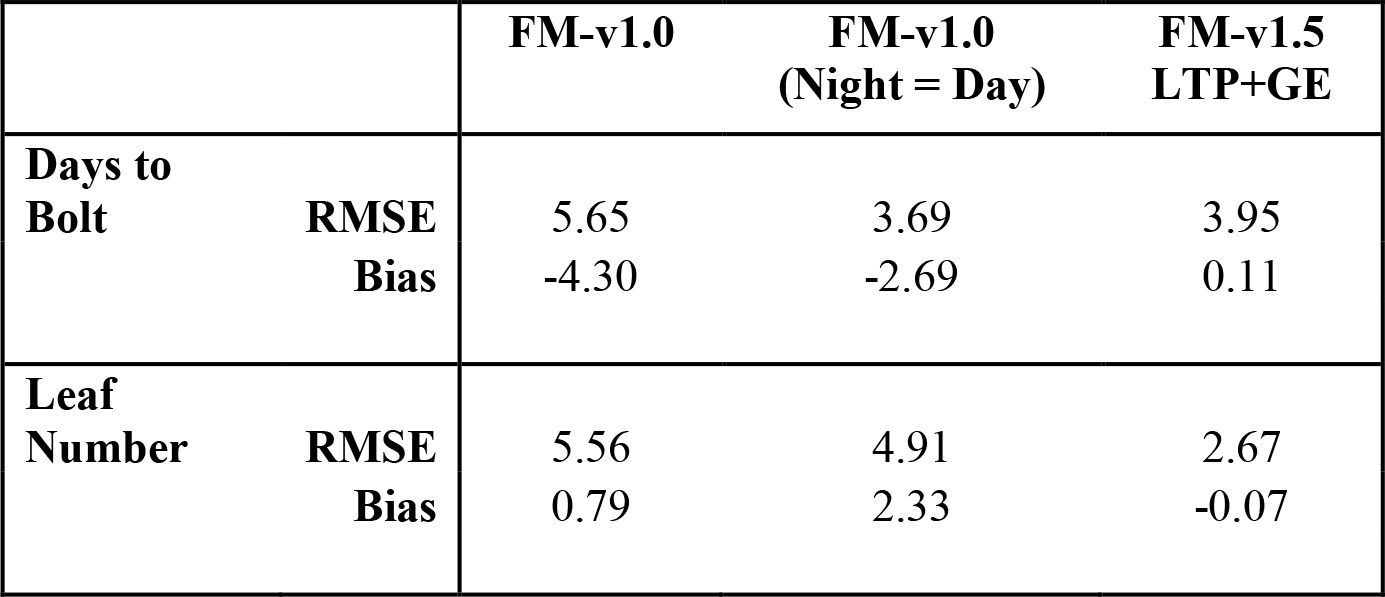
Fit of FM-v1.0 and FM-v1.5 for Columbia-0 and Landsberg *erecta* combined.

### *Influence of* FT *accumulation in conditions causing later flowering*

As later produced leaves may lose the capacity to express *FT* (Figure 2), we wondered how this would impact *FT* accumulation and flowering over longer developmental time periods, such as in cool constant temperatures when *FT* is suppressed and *Arabidopsis* flowers at a higher leaf number (Blazquez *et al.*, 2003). We grew Col-0 at 12 °C-constant or 22/12°C-day conditions (in the latter treatment, plants then remained at 12°C). We observed flowering at 24 and 28 leaves, respectively, and at 60 and 61 days after sowing. In the full FM-v1.5 LTP+GE variant, *FT* failed to accumulate to the threshold set in 22 °C conditions (Figure 6). Simulated *FT* in the LTP variant (temperature influence on *FT* gene expression removed), did reach the threshold in 22°C/12°C-day conditions (data not shown). *FT* attained the threshold in 12°C-constant, only after influence of leaf age was removed from the LTP model as well. Therefore, whole-plant *FT* accumulation, as influenced by leaf age, leaf tissue production, and transcriptional regulation of *FT* by temperature may not be sufficient to predict flowering in conditions in which *FT* is strongly suppressed under the assumption of a constant *FT* threshold.

**Figure 6:**
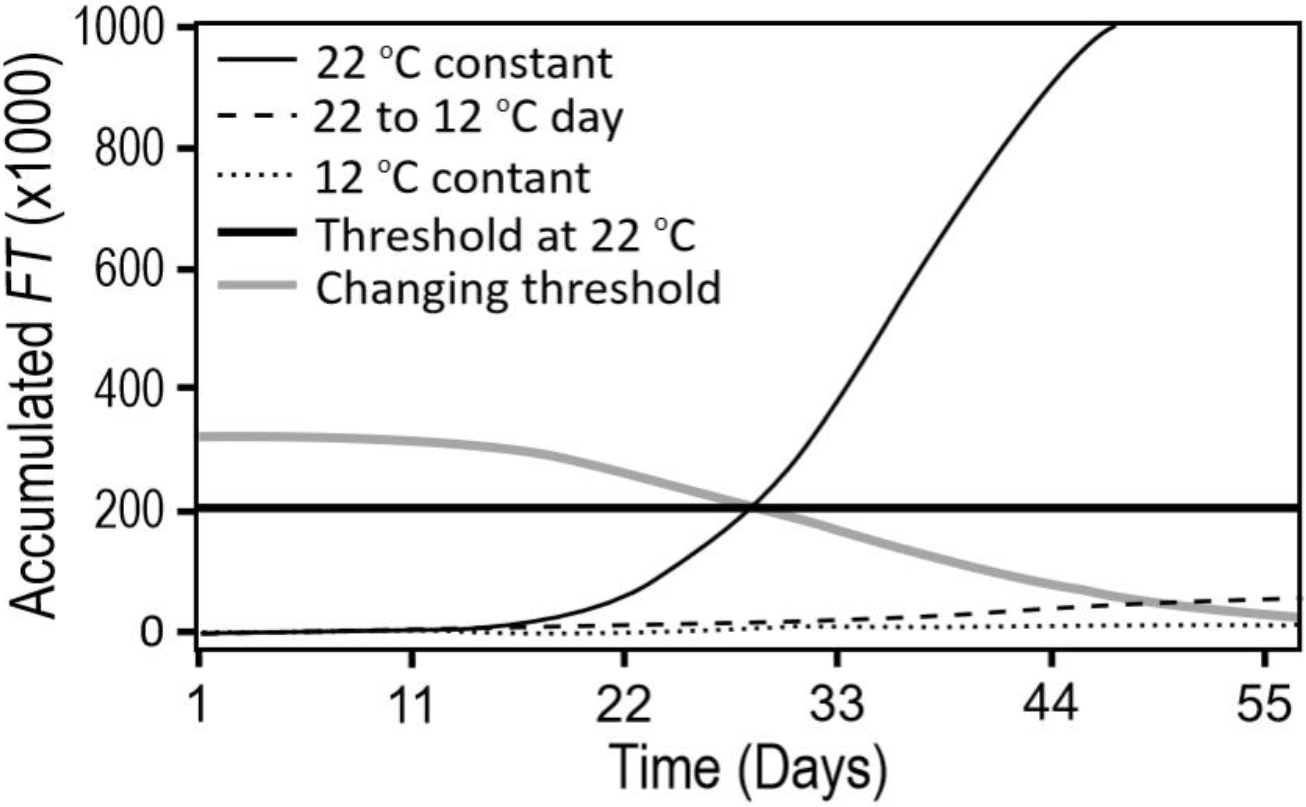
Plants grown at constant cool (12 °C) temperatures from seed (constant) or after one week at 22 °C (22 to 12 °C day) do not accumulate *FT* to a threshold set using 22 °C constant temperatures in long days (thick black line). Altering the threshold to decline with developmental time (thick gray line) improves the predictive capacity of FM-v1.5.

### *Influence of short-term temperature fluctuations on* FT *and flowering*

Although long-term exposure to cool temperatures suppressed whole-plant *FT* and delayed flowering, temperature changes at dawn in LDs (22/12°C-day or 22/17°C-day) caused short-term *FT* induction (Kinmonth-Schultz et al. 2016). As *FT* transcript must accumulate over several days before flowering can occur (Krzymuski *et al.*, 2015), we wondered whether a short-term temperature drop, causing *FT* induction, could complement *FT* produced in subsequent warm temperatures to accelerate flowering, or if slower whole-plant *FT* accumulation with slower leaf growth would delay flowering. To compare the predicted influence of *FT* induction by temperature fluctuations, we used the FM-v1.5 LTP+GE and LTP variants to simulate two-week-old plants moved to 12 °C in LDs for two, four, or six days (12°C-2d, -4d, or -6d), then moved to warm, LD conditions. We also grew plants in these conditions. Control plants were moved directly to warm, LD conditions at two weeks.

Simulating these conditions in the full LTP+GE variant of FM-v1.5, we found little difference in days to bolt between 12°C-2d and the control and a three-day difference between 12°C-6d and the control. There was a decline in leaf number from 15 to 14 leaves in plants exposed to 12°C-2d and 12°C-4d, indicating flowering at a slightly younger developmental age that translated to little difference in days to bolt between the control and 12°C-2d. In 12°C-6d, the leaf number increased again to be like the control. In the LTP variant, the leaf number of all three treatments was the same as the control, whereas there was an increase in days to bolt for each consecutive two-days at 12 °C, consistent with slowed accumulation of *FT* due to slower leaf growth.

We observed slowed growth (relative to the control) in the cool-temperature treatments. Visible leaf number was significantly lower after four and six days in 12 °C (Figure 7a). On day seven, after completion of all cool-temperature treatments, there was a gradient in leaf area across treatments, with plants from 12°C-6d being the smallest (Figure 7b, S8). We observed a statistically significant delay in the number of days to visible bolt in both 12°C-4d and12°C-6d, like both simulations (*P*<0.001, Table 3, Figure 7c). While we did not observe a significant difference in leaf number in either 12°C-2d or 12°C-4d relative to the control, plants in 12°C-6d produced approximately 1.5 more leaves before flowering than the other three treatments (*P*<0.001), more like the predicted increase in leaf number from 12°C-2d and 12°C-4d to 12°C-6d in the LTP+GE model variant (Table 3).

**Figure 7:**
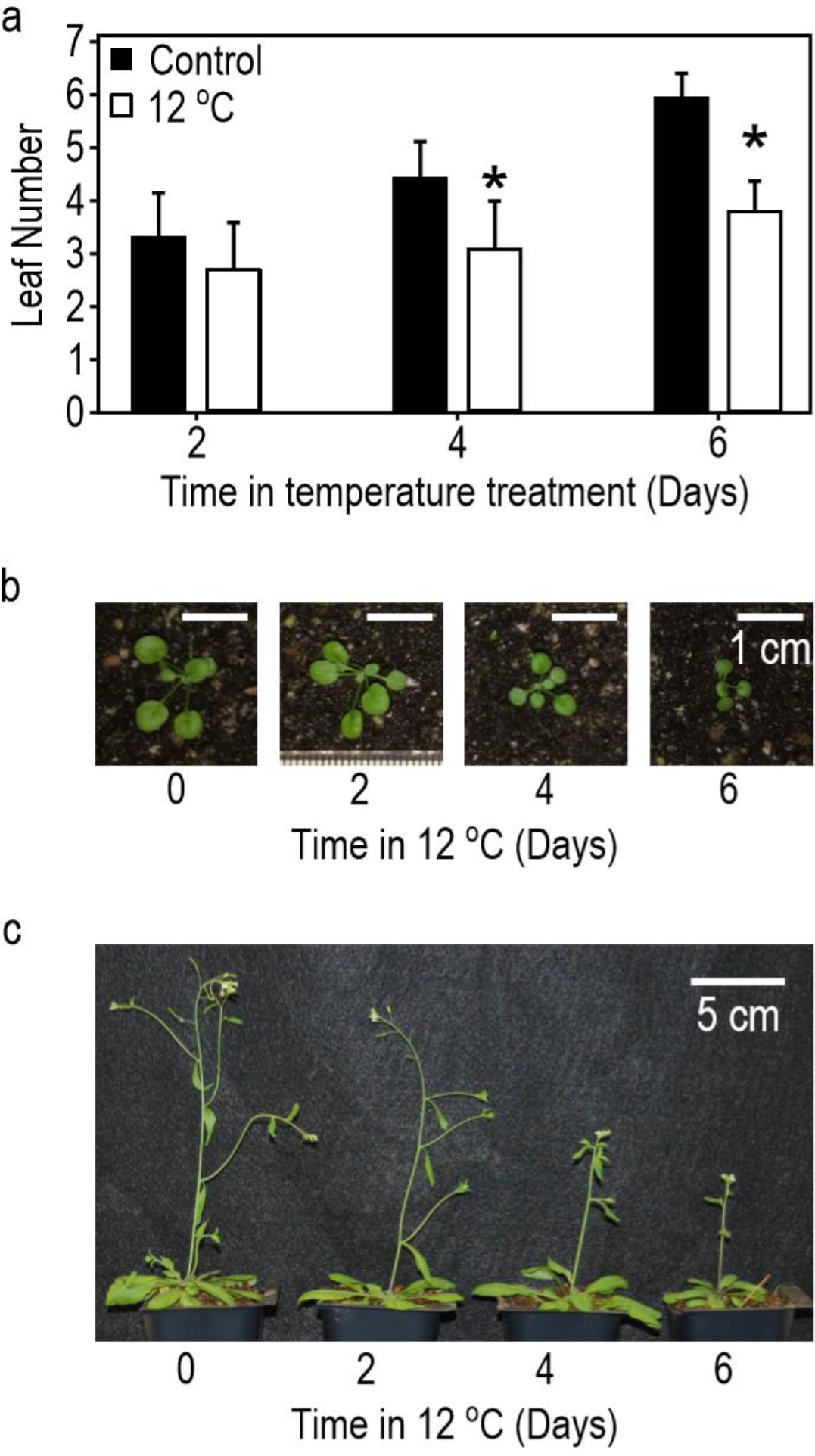
Growth is slowed and flowering is delayed in plants exposed to 12 °C for two, four, or six days, then returned to warm temperatures (24 °C), relative to control plants grown continuously in warm-temperatures. **(a)** Average leaf number of plants recorded at dawn after two, four, or six days in 24 °C (control) or 12 °C temperature conditions. **(b)** Relative seedling sizes on dawn of day seven, after completion of all cool-temperature treatments (scale bars = 1cm, 0 = control). Individual images cropped from the same photograph and scaled together (see original image, Figure S9). **(c)** Relative flowering progression three days after appearance of last floral stem (bolt) in plants exposed to 12 °C for two, four, or six days relative to 24 °C control (0, scale bar = 5cm).

**Table 3:** Observed and simulated days to bolt and leaf number of rosette leaves on the main stem in Columbia plants exposed to short-term drops 12 °C temperature relative to plants remaining in the warm temperature control (24 °C) in long days (LD).

|  | treatment | Obs.<br>Data | n | Robust<br>S.E. | Robust<br>z | P-Value | C.I. of<br>dif.<br>(lower) | C.I. of<br>dif.<br>(upper) | FM-v1.5<br>LTP+GE | FM-v1.5<br>LTP |
| --- | --- | --- | --- | --- | --- | --- | --- | --- | --- | --- |
| **Days to bolt** | LD24C<br>(int) | 35.00 | 11 | 0.62 | 31.10 |  |  |  | 34.75 | 34.75 |
|  | 12C, 2d | 36.00 | 15 | 0.60 | 1.73 | 0.08 | -0.65 | 2.73 | 34.92 | 35.75 |
|  | 12C, 4d | 37.31 | 14 | 0.42 | 5.54 | 0.00 | 0.87 | 3.80 | 35.83 | 36.62 |
|  | 12C, 6d | 38.64 | 14 | 0.35 | 7.85 | 0.00 | 2.24 | 5.02 | 37.79 | 37.58 |
| **Leaf**<br>**number** | LD24C<br>(int) | 13.73 | 11 | 0.52 | 19.59 |  |  |  | 15.00 | 15.00 |
|  | 12C, 2d | 13.13 | 15 | 0.34 | -1.21 | 0.23 | -0.81 | 1.65 | 14.00 | 15.00 |
|  | 12C, 4d | 13.64 | 14 | 0.36 | 0.49 | 0.63 | -1.07 | 1.42 | 14.00 | 15.00 |
|  | 12C, 6d | 15.14 | 14 | 0.39 | 3.47 | 0.00 | 0.08 | 2.64 | 15.00 | 15.00 |

Observed treatments counted significantly different from the control when *P*<0.05 and the confidence interval of the difference from the control does not contain zero.

## Discussion

Incorporating underlying mechanisms could improve model utility for a range of conditions without requiring recalibration (White, 2009; Boote *et al.*, 2013). Here, we found that thermal time (MPTUs) did predict delays in days to bolt under fluctuating temperature conditions in LDs relative to the constant-temperature control, but the delays were less than observed and more like FM-v1.5 LTP, in which *FT* accumulated only with leaf growth, a function of thermal time (Table 1, Figure 5b & d). Adding direct temperature regulation of *FT* improved model predictions by increasing the degree of predicted difference between the warm-day, cool-night treatments and the control.

*FT* was reduced in later-produced leaves (Figure 2). This change in *FT* expression with developmental age was incorporated into FM-v1.5 using leaf age as a proxy, and caused *FT* to fail to accumulate to a preset threshold to predict flowering in constant cool temperatures. This finding enables integration of qualitative (presence/absence) and quantitative (dosage response) aspects of *FT* effects on flowering, and has implications for other conditions in which *FT* is suppressed, such as in short daylengths. It can help us quantify when *FT* plays a role during development, when *FT* alone is a poor predictor of flowering, and when it may act synergistically or competitively with other flowering factors.

For instance, the *FT* threshold requirement should be influenced by shoot-apex genes; their sensitivity likely changes with climate and developmental age. For example, in short-days, high temperatures may reduce SVP activity at the shoot apex to initiate flowering despite lower *FT* levels (Fernández *et al.*, 2016). At the shoot apex, SVP suppresses *SUPPRESSOR OF OVEREXPRESSION OF CONSTANS* (*SOC1*), which is positively regulated by *FT*, and which activates *LEAFY* (*LFY*), a key player in the floral transition (Schmid *et al.*, 2003; Lee *et al.*, 2008; Jang *et al.*, 2009). FT protein also activates *APETALA1* (*AP1*) at the shoot apex (Lee & Lee, 2010). AP1, in turn, is involved in the down regulation of *TERMINAL FLOWERING1* (*TFL1*), a *FT* homolog. TFL1 is thought to compete with FT for binding with FD to suppress *LFY*, as well as *AP1*, forming a negative feedback loop (Kaufmann *et al.*, 2010; Wickland & Hanzawa, 2015). Both the decrease in SVP and TFL1 would likely decrease the *FT* threshold needed to induce flowering. Like *SVP, TFL1* may be temperature sensitive (Kim *et al.*, 2013).

A changing threshold, due to different *LATE FLOWERING* alleles in Pea, a homologue of *TFL1* in *Arabidopsis* (Foucher *et al.*, 2003), aids flowering time predictions (Wenden *et al.*, 2009). Incorporating such a mechanism – influenced by climate and developmental age – may aid understanding of how climate influences flowering. As proof of concept, we caused the *FT* threshold level to change with developmental age (thermal time) (Figure 6). Doing so improved the predictive capacity of FM-v1.5 in constant, cool temperatures.

SVP, in conjunction with FLM, suppresses *FT* in response to cool temperatures (Blazquez *et al.*, 2003; Lee *et al.*, 2007, 2013). We demonstrated that residual SVP and FLM activity after shortterm cold exposures could be important for *FT* regulation. For instance, to mimic observed dusk suppression of *FT* in warm-day, cool-night temperature cycles, simulated SVP activity decayed slowly after at 12 °C night, such that it was higher after 16 hs at 22 °C, than it was in constant 22 °C conditions. Our model also highlights the need to clarify the degree of temperature influence in *FT* activation and suppression at a range of temperatures. For example, in FM-v1.5, *FT* is not induced to observed levels, and induction is not maintained as long, after dawn exposure to 17 °C (Figure 3f). It is possible that SVP activation is lower in 17 °C, than predicted from our model. However, the relative difference in transcript levels across treatments is similar to the relative difference in daytime *FT* expression, which correlates most strongly with flowering (Krzymuski *et al.*, 2015; Kinmonth-Schultz *et al.*, 2016).

Our simulations, while requiring validation in other temperature conditions, are consistent with approaches that use day length and vernalization to influence the leaf number at which the reproductive transition occurs (Brown *et al.*, 2013). However, our work demonstrates that ambient temperature should be incorporated to influence leaf number as well, not only developmental rate. For instance, we altered *FT* accumulation, either by removing temperature influence on *FT* transcription (FM-v1.5 LTP, Table 2) or by short-term, cool-temperature exposure (Figure 3d-e, Table 3), affecting final leaf number. In each instance, *FT* still accumulated with leaf production as influenced by temperature, demonstrating that temperature influences when (in days) the reproductive transition occurs by influencing the developmental rate and whole-plant *FT* accumulation. We further suggest that tissue accumulation through growth is an underlying factor in the accumulation of thermal time as it causes gene products to accumulate. Together, this work demonstrates that decomposing the influences of climate and development can improve our understanding of plant responses in a range of conditions.

## Supplementary Data

**Section S1:** Materials and Methods for plant growth conditions, RNA expression, GUS tissue analysis, and statistical analysis and experimental controls.

**Section S2:** Equation used in FM-v1.0 to describe *FT* transcription as a function of CO protein.

**Table S1:** Coefficients values for equations used in FM-v1.5.

**Figure S1.** Graphic representation of FM-V1.

**Figure S2.** SVP/FLM activity declines over time.

**Figure S3.** Behavior of *CO* mRNA in response to different temperature regimes.

**Figure S4.** Simulated expression profile of *LHY*, plotted over time used to increase morning transcriptional sensitivity of *FT*.

**Figure S5.** The spatial expression profile of *FT* changes with leaf age.

**Figure S6:** Behavior of morphological and physiological parameters in FM-v1.0 and v1.5.

**Figure S7.** Original photograph used for Figure 7 showing Specific Leaf Area (SLA) declines after growth in cool constant temperatures or in warm-day, cool-night temperature cycles relative to a constant, warm-temperature control.

**Figure S8.** Simulated *FT* expression profile in FM-v1.5 in the *svp* mutant mimics the pattern but not relative amplitude of that observed.

## Acknowledgements

Many thanks the Jennifer Nemhauser lab for use of their dissecting scope and to Paul Panipinto for help on this project. This work was supported in part by a Cooperative Research Program for Agricultural Science and Technology Development (PJ0127872017), Rural Development Administration, Republic of Korea and a Specific Cooperative Agreement (58-8042-6-097) between University of Washington and USDA-ARS to S.-H.K; by National Institute of Health grant (GM079712), Next-Generation BioGreen 21 Program (SSAC, PJ011175, Rural Development Administration, Republic of Korea) to T.I.; by a Biotechnology and Biological Sciences Research Council awards (BB/L026996, BB/N012348) to A.J.M.; and by the University of Washington Biology Department Frye-Hotson-Rigg Fellowship to H.K.S.

## References

Abe M, Kobayashi Y, Yamamoto S, Daimon Y, Yamaguchi A, Ikeda Y, Ichinoki H, Notaguchi M, Goto K, Araki T. 2005. FD, a bZIP protein mediating signals from the floral pathway integrator FT at the shoot apex. Science 309: 1052–6.

Amasino R. 2010. Seasonal and developmental timing of flowering. Plant Journal 61: 1001–1013.

Asseng S, Ewert F, Rosenzweig C, Jones JW, Hatfield JL, Ruane AC, Boote KJ, Thorburn PJ, Rötter RP, Cammarano D, et al. 2013. Uncertainty in simulating wheat yields under climate change. Nature Climate Change 3: 827–832.

Blazquez M, Ahn J, Weigel D. 2003. A thermosensory pathway controlling flowering time in Arabidopsis thaliana. Nature Genetics 33: 168–171.

Boote KJ, Jones JW, White JW, Asseng S, Lizaso JI. 2013. Putting mechanisms into crop production models. Plant, Cell and Environment 36: 1658–1672.

Bratzel F, Turck F. 2015. Molecular memories in the regulation of seasonal flowering: from competence to cessation. Genome Biology 16: 192–206.

Brown HE, Jamieson PD, Brooking IR, Moot DJ, Huth NI. 2013. Integration of molecular and physiological models to explain time of anthesis in wheat. Annals of Botany 112: 1683–1703.

Carter JM, Orive ME, Gerhart LM, Stern JH, Marchin RM, Nagel J, Ward JK. 2017. Warmest extreme year in U.S. history alters thermal requirements for tree phenology. Oecologia 183: 1197–1210.

Chew YH, Wenden B, Flis A, Mengin V, Taylor J, Davey CL, Tindal C, Thomas H, Ougham HJ, Reffye P de, et al. 2014. Multiscale digital Arabidopsis predicts individual organ and whole-organism growth. Proceedings of the National Academy of Sciences 111: E4127–E4136.

Chew YH, Wilczek AM, Williams M, Welch SM, Schmitt J, Halliday KJ. 2012. An augmented Arabidopsis phenology model reveals seasonal temperature control of flowering time. New Phytologist 194: 654–665.

Christophe A, Letort V, Hummel I, Cournède P-H, Reffye P de, Lecœur J. 2008. A model-based analysis of the dynamics of carbon balance at the whole-plant level in Arabidopsis thaliana. Functional Plant Biology 35: 1147–1162.

Chuine I. 2000. A unified model for budburst of trees. Journal of Theoretical Biology 207: 337–347.

Corbesier L, Vincent C, Jang S, Fornara F, Fan Q, Searle I, Giakountis A, Farrona S, Gissot L, Turnbull C, et al. 2007. FT protein movement contributes to long-distance signaling in floral induction of Arabidopsis thaliana. Science 316: 1030–1033.

Fernández V, Takahashi Y, LeGourrierec J, Coupland G. 2016. Photoperiodic and thermosensory pathways interact through CONSTANS to promote flowering at high temperature under short days. Plant Journal 86: 426–440.

Foucher F, Morin J, Courtiade J, Cadioux S, Ellis N, Banfield MJ, Rameau C. 2003. DETERMINATE and LATE FLOWERING are two TERMINAL FLOWER1/CENTRORADIALIS homologs that control two distinct phases of flowering initiation and development in pea. Plant Cell 15: 2742–2754.

Franklin KA, Whitelam GC. 2007. Light-quality regulation of freezing tolerance in Arabidopsis thaliana. Nat Genet 39:1410–1413.

He J, Le Gouis J, Stratonovitch P, Allard V, Gaju O, Heumez E, Orford S, Griffiths S, Snape JW, Foulkes MJ, et al. 2012. Simulation of environmental and genotypic variations of final leaf number and anthesis date for wheat. European Journal of Agronomy 42: 22–33.

Heinsvig Kjær K, Thorup-Kristensen K, Rosenqvist E, Mazanti Aaslyng J. 2007. Low night temperatures change whole-plant physiology and increase starch accumulation in Chrysanthemum morifolium. Journal of Horticultural Science & Biotechnology 82: 867–874.

Helliwell CA, Wood CC, Robertson M, James Peacock W, Dennis ES. 2006. The Arabidopsis thaliana FLC protein interacts directly in vivo with SOC1 and FT chromatin and is part of a high-molecular-weight protein complex. Plant Journal 46: 183–192.

Jaglo-Ottosen KR, Gilmour SJ, Zarka DG, Schabenberger O, Thomashow MF. 1998. Arabidopsis CBF1 overexpression induces COR genes and enhances freezing tolerance. Science 280: 104–106.

Jamieson PD, Brooking IR, Semenov MA, Porter JR. 1998a. Making sense of wheat development: a critique of methodology. Field Crops Research 55: 117–127.

Jamieson PD, Semenov MA, Brooking IR, Francis GS. 1998b. Sirius: a mechanistic model of wheat response to environmental variation. European Journal of Agronomy 8: 161–179.

Jang S, Torti S, Coupland G. 2009. Genetic and spatial interactions between FT, TSF and SVP during the early stages of floral induction in Arabidopsis. Plant Journal 60: 614–625.

Jones JW, Hoogenboom G, Porter CH, Boote KJ, Batchelor WD, Hunt LA, Wilkens PW, Singh U, Gijsman AJ, Ritchie JT. 2003. The DSSAT cropping system model. European Journal of Agronomy 18: 235–265.

Karsai I, Igartua E, Casas AM, Kiss T, Soos V, Balla K, Bedo Z, Veisz O. 2013. Developmental patterns of a large set of barley (Hordeum vulgare) cultivars in response to ambient temperature. Annals of Applied Biology 162: 309–323.

Karsai I, Szucs P, Koszegi B, Hayes PM, Casas A, Bedo Z, Veisz O. 2008. Effects of photo and thermo cycles on flowering time in barley: a genetical phenomics approach. Journal of Experimental Botany 59: 2707–2715.

Kaufmann K, Wellmer F, Muiño JM, Ferrier T, Wuest SE, Kumar V, Serrano-Mislata A, Madueño F, Krajewski P, Meyerowitz EM, et al. 2010. Orchestration of floral initiation by APETALA1. Science 328: 85–89.

Kim S-H, Gitz DC, Sicher RC, Baker JT, Timlin DJ, Reddy VR. 2007. Temperature dependence of growth, development, and photosynthesis in maze under elevated CO_2_. Environmental and Experimental Botany 61: 224–236.

Kim W, Park TI, Yoo SJ, Jun AR, Ahn JH. 2013. Generation and analysis of a complete mutant set for the Arabidopsis FT/TFL1 family shows specific effects on thermo-sensitive flowering regulation. Journal of Experimental Botany 64: 1715–1729.

Kim S-H, Yang Y, Timlin DJ, Fleisher DH, Dathe A, Reddy VR, Staver K. 2012. Modeling temperature responses of leaf growth, development, and biomass in maize with MAIZSIM. Agronomy Journal 104: 1523–1537.

Kinmonth-Schultz HA, Tong X, Lee J, Song YH, Ito S, Kim S-H, Imaizumi T. 2016. Cool night-time temperatures induce the expression of CONSTANS and FLOWERING LOCUS Tto regulate flowering in Arabidopsis. New Phytologist 211: 208–224.

Krzymuski M, Andrés F, Cagnola JI, Seonghoe J, Yanovsky M, Coupland G, Casal JJ. 2015. The dynamics of FLOWERING LOCUS T expression encodes long-day information. Plant journal 83: 952–961.

Kumudini S, Andrade FH, Boote KJ, Brown GA, Dzotsi KA, Edmeades GO, Gocken T, Goodwin M, Halter AL, Hammer GL, et al. 2014. Predicting maize phenology: Intercomparison of functions for developmental response to temperature. Agronomy Journal 106: 2087–2097.

Lee J, Lee I. 2010. Regulation and function of SOC1, a flowering pathway integrator. Journal of Experimental Botany 61: 2247–2254.

Lee J, Oh M, Park H, Lee I. 2008. SOC1 translocated to the nucleus by interaction with AGL24 directly regulates LEAFY. Plant Journal 55: 832–843.

Lee JH, Ryu H-S, Chung KS, Posé D, Kim S, Schmid M, Ahn JH. 2013. Regulation of temperature-responsive flowering by MADS-Box transcription factor repressors. Science 342: 628–632.

Lee JH, Yoo SJ, Park SH, Hwang I, Lee JS, Ahn JH. 2007. Role of SVP in the control of flowering time by ambient temperature in Arabidopsis. Genes & Development 21: 397–402.

Lehenbauer PA. 1914. Growth of maize seedlings in relation to temperature. Physiological Researches 1: 247–288.

Lutz U, Posé D, Pfeifer M, Gundlach H, Hagmann J, Wang C, Weigel D, Mayer KFX, Schmid M, Schwechheimer C. 2015. Modulation of ambient temperature-dependent flowering in Arabidopsis thaliana by natural variation of FLOWERING LOCUS M. PLoS Genetics 11: e1005588.

Makowski D, Asseng S, Ewert F, Bassu S, Durand JL, Li T, Martre P, Adam M, Aggarwal PK, Angulo C, et al. 2015. A statistical analysis of three ensembles of crop model responses to temperature and CO_2_ concentration. Agricultural and Forest Meteorology 214: 483–493.

Mathieu J, Yant LJ, Mürdter F, Küttner F, Schmid M. 2009. Repression of flowering by the miR172 target SMZ. PLoS Biology 7: e1000148.

Parent B, Turc O, Gibon Y, Stitt M, Tardieu F. 2010. Modelling temperature-compensated physiological rates, based on the co-ordination of responses to temperature of developmental processes. Journal of Experimental Botany 61: 2057–2069.

Piper EL, Smit MA, Boote KJ, Jones JW. 1996. The role of daily minimum temperature in modulating the development rate to flowering in soybean. Field Crops Research 47: 211–220.

Posé D, Verhage L, Ott F, Yant L, Mathieu J, Angenent GC, Immink RGH, Schmid M. 2013. Temperature-dependent regulation of flowering by antagonistic FLM variants. Nature 503: 414–417.

Pyl E-T, Piques M, Ivakov A, Schulze W, Ishihara H, Stitt M, Sulpice R. 2012. Metabolism and growth in Arabidopsis depend on the daytime temperature but are temperature-compensated against cool nights. Plant Cell 24: 2443–2469.

Rasse DP, Tocquin P. 2006. Leaf carbohydrate controls over Arabidopsis growth and response to elevated CO_2_: an experimentally based model. New Phytologist 172: 500–513.

Rensing L, Ruoff P. 2002. Temperature effect on entrainment, phase shifting, and amplitude of circadian clocks and its molecular bases. Chronobiology International 19: 807–864.

Ritchie J, Otter S. 1985. Description and performance of CERES-Wheat. A user-oriented wheat model. US Department of Agriculture, ARS 38: 159–175.

Salazar JD, Saithong T, Brown PE, Foreman J, Locke JCW, Halliday KJ, Carré IA, Rand DA, Millar AJ. 2009. Prediction of photoperiodic regulators from quantitative gene circuit models. Cell 139: 1170–1179.

Schmid M, Uhlenhaut NH, Godard F, Demar M, Bressan R, Weigel D, Lohmann JU. 2003. Dissection of floral induction pathways using global expression analysis. Development 130: 6001–6012.

Schwartz C, Balasubramanian S, Warthmann N, Michael TP, Lempe J, Sureshkumar S, Kobayashi Y, Maloof JN, Borevitz JO, Chory J, et al. 2009. Cis-regulatory changes at FLOWERING LOCUS T mediate natural variation in flowering responses of Arabidopsis thaliana. Genetics 183: 723–732.

Searle I, He Y, Turck F, Vincent C, Fornara F, Kröber S, Amasino RA, Coupland G. 2006. The transcription factor FLC confers a flowering response to vernalization by repressing meristem competence and systemic signaling in Arabidopsis. Genes & Development 20: 898–912.

Seaton DD, Smith RW, Song YH, MacGregor DR, Stewart K, Steel G, Foreman J, Penfield S, Imaizumi T, Millar AJ, et al. 2015. Linked circadian outputs control elongation growth and flowering in response to photoperiod and temperature. Molecular Systems Biology 11: 1–19.

Segal IH. 1976. Biochemical Calculations: How to Solve Mathematical Problems in General Biochemistry. Canada: John Wiley and Sons.

Song Y-H, Shim J-S, Kinmonth-Schultz HA, Imaizumi T. 2015. Photoperiodic flowering: time measurement mechanisms in leaves. Annual Review of Plant Biology 66: 441–464.

Sperling O, Earles JM, Secchi F, Godfrey J, Zwieniecki MA. 2015. Frost induces respiration and accelerates carbon depletion in trees. Plos One 10: e0144124.

Sureshkumar S, Dent C, Seleznev A, Tasset C, Balasubramanian S. 2016. Nonsense-mediated mRNA decay modulates FLM-dependent thermosensory flowering response in Arabidopsis. Nature Plants 2: 16055.

Takada S, Goto K. 2003. TERMINAL FLOWER2, an Arabidopsis homolog of HETEROCHROMATIN PROTEIN1, counteracts the activation of FLOWERING LOCUS Tby CONSTANS in the vascular tissues of leaves to regulate flowering time. Plant Cell 15: 2856–2865.

Thines BC, Youn Y, Duarte MI, Harmon FG. 2014. The time of day effects of warm temperature on flowering time involve PIF4 and PIF5. Journal of Experimental Botany 65: 1141–1151.

Welch SM, Roe JL, Dong Z. 2003. A genetic neural network model of flowering time control in Arabidopsis thaliana. Agronomy Journal 95: 71–81.

Wenden B, Dun EA, Hanan J, Andrieu B, Weller JL, Beveridge CA, Rameau C. 2009. Computational analysis of flowering in pea (Pisum sativum). New Phytologist 184: 153–167.

White JW. 2009. Combining ecophysiological models and genomics to decipher the GEM-to-P problem. NJAS - Wageningen Journal of Life Sciences 57: 53–58.

Wickland DP, Hanzawa Y. 2015. The FLOWERING LOCUS T/TERMINAL FLOWER 1 gene family: functional evolution and molecular mechanisms. Molecular Plant 8: 983–997.

Wilczek AM, Roe JL, Knapp MC, Cooper MD, Lopez-Gallego C, Martin LJ, Muir CD, Sim S, Walker A, Anderson J, et al. 2009. Effects of genetic perturbation on seasonal life history plasticity. Science 323: 930–934.

Yan L, Fu D, Li C, Blechl A, Tranquilli G, Bonafede M, Sanchez A, Valarik M, Yasuda S, Dubcovsky J. 2006. The wheat and barley vernalization gene VRN3 is an orthologue of FT. Proceedings of the National Academy of Sciences 103: 19581–19586.

Yin X, Kropff M. 1996. The effect of temperature on leaf appearance in rice. Annals of Botany 77: 215–221.

Yin X, Kropff MJ, McLaren G, Visperas RM. 1995. A nonlinear model for crop development as a function of temperature. Agricultural and Forest Meteorology 77: 1–16.

Zheng B, Biddulph B, Li D, Kuchel H, Chapman S. 2013. Quantification of the effects of VRN1 and Ppd-D1 to predict spring wheat (Triticum aestivum) heading time across diverse environments. Journal of Experimental Botany 64: 3747–3761.

